# A novel method for obtaining epileptic brain tissue for omic analyses using electrodes from a clinical stereoelectroencephalography study

**DOI:** 10.64898/2026.09.01.747265

**Authors:** Ninja Ehrnrooth, Salla M. Kangas, Alina Malyutina, Juha Wilenius, Antti Tuhkala, Salla Keskitalo, Kari Salokas, Markku Varjosalo, Hanna Lindgren, Anne Mäkelä, Olli Tynninen, Atte Karppinen, Leena Lauronen, Jukka Vanhanen, Päivi Nevalainen, Maria Peltola, Johanna Uusimaa, Reetta Hinttala, Eeva-Liisa Metsähonkala, Erika Ignatius

## Abstract

**Purpose:** Developing a valid and practical method for obtaining brain tissue samples for transcriptomic and proteomic analyses using tissue adhered to stereoelectroencephalography (SEEG) electrodes, to enable investigation of the molecular mechanisms underlying chronic epilepsy.

**Method:** Brain tissue samples adhered to SEEG electrodes were collected from six patients with drug-resistant epilepsy and preprocessed in a hospital environment after electrode removal. RNA extraction was initiated immediately, and proteomics samples were snapfrozen until subsequent analysis.

The samples were categorized into three groups according to their electrophysiological profile: epileptogenic zone, propagation zone, and least-involved zone. The omic findings between these zones and anatomical brain areas were compared.

**Results:** High-quality RNA and protein samples were obtained from tissue adhered to SEEG electrodes. Neuron- and brain-specific gene expression patterns and proteins were identified. Signs of activation of inflammatory mechanisms were most pronounced in the epileptogenic zone. Transcriptomic and proteomic findings demonstrated concordance.

**Conclusion:** SEEG electrodes are a useful source for obtaining brain tissue for molecular characterization of chronic epilepsy. This method will enable the identification of shared and distinct molecular mechanisms in patients with varying etiologies of epilepsy.

## 1 Introduction

Despite substantial advances in epilepsy treatment possibilities and diagnostic techniques, severe epilepsy remains a major clinical challenge. Approximately one third of patients continue to have seizures, despite major advances in epilepsy management (Kwan et al., 2010). Beyond the immediate effects of recurrent seizures, severe epilepsy can have profound consequences for cognitive function, psychosocial well-being, and quality of life, and may disrupt the normal development and organization of neuronal networks (Fisher et al., 2005).

Early diagnosis and intervention may help prevent the progression of epilepsy to a severe and drug-resistant epilepsy (Tchaicha et al., 2026). However, achieving this goal requires a better understanding of the molecular and cellular processes that drive epileptogenesis and shape epileptic networks. Investigating how these biological mechanisms interact and evolve over time could reveal new opportunities for disease modification and the development of more effective therapies.

Different omic studies have been conducted to investigate the biology of epilepsy using animal models, cerebrospinal fluid samples, human epilepsy surgery resection samples, and human postmortem samples. While each provides valuable knowledge, all are associated with important limitations.

In patients with drug-resistant focal epilepsy, stereoelectroencephalography (SEEG) is used to localize and delineate epileptogenic networks and guide surgical treatment (Isnard et al., 2018). During this procedure, depth electrodes are implanted into suspected epileptogenic brain regions and later removed after the monitoring period. Small amounts of brain tissue remain attached to the surfaces of the electrodes.

Recent studies have demonstrated that these tissue remnants represent a valuable source of molecular information. DNA extracted from SEEG electrodes has been successfully used to identify somatic genetic variants associated with epilepsy (Checri et al., 2023; Montier et al., 2019; Ye et al., 2022) and more recent reports have shown that transcriptomic profiling is also feasible using this material (Dwivedi et al., 2025; Larkin et al., 2025). These findings highlight the unique potential of SEEG-derived tissue to provide molecular information from precisely defined regions of the living human epileptic brain.

In this study, we sought to develop a practical and easily implementable workflow for collecting brain tissue from SEEG electrodes for both transcriptomic and proteomic analyses. The protocol was adapted from a previously described method developed for tissue derived from guide tubes and recording electrodes used in routine deep brain stimulation implantation procedures (Kangas et al., 2022), with the aim of making it readily applicable in routine clinical practice. Using this approach, we generated transcriptomic and proteomic datasets from SEEG-derived brain tissue and characterized their molecular features. We also present preliminary findings demonstrating molecular differences between brain regions exhibiting distinct levels of epileptic activity.

## 2 Methods

### 2.1 Patients

SEEG implantations and recordings were performed as part of epilepsy surgery evaluation at Helsinki University Hospital, Helsinki, Finland. All six patients with drug-resistant epilepsy (DRE) who underwent a SEEG examination between February and June 2024 provided informed consent to participate in the study. Samples for transcriptomic and proteomic analyses were collected from five patients (two adult and three pediatric patients); an additional pediatric patient contributed samples for only proteomic analyses in this study. The clinical characteristics of the study cohort are summarized in Tables 1.

**Table 1.**
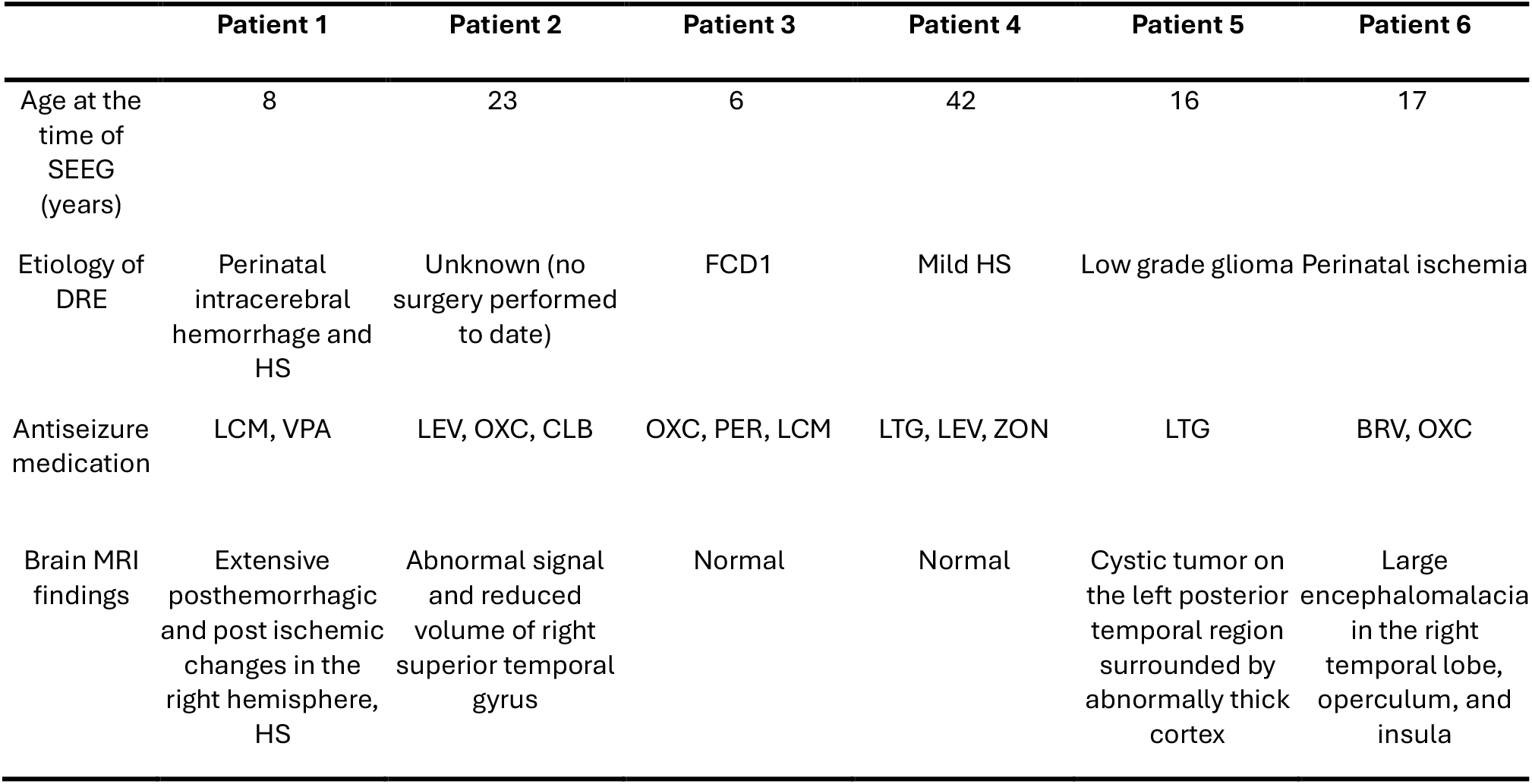
Clinical characteristics of the patients. Abbreviations: SEEG, stereoelectroencephalOgraphy; BRV, brivaracetam; CLB; clobazam; DRE, drug-resistant epilepsy; FCD, focal cortical dysplasia; HS hippocampal sclerosis; LCM, lacosamide; LEV, levetiracetam; LTG, lamotrigine; MRI, magnetic resonance imaging; OXC, oxcarbazepine; PER, perampanel; VPA, valproate; ZON, zonisamide

### 2.2 SEEG recordings

The SEEG implantations and recordings were performed according to current clinical guidelines (Isnard et al., 2018). SEEG electrodes from DIXI Medical (DIXI Medical, Marchaux - Chaudefontain, France) were implanted using either a Leksell (Elekta AB, Stockholm, Sweden) stereotactic frame or a Neuromate® stereotactic robot (Renishaw plc, Gloucestershire, UK). Postoperatively, the precise location of each implanted electrode was confirmed by co-registration of postoperative CT images with preoperative anatomical MRI scans. SEEG recordings with a duration of 7-9 days were done using the Micromed SD LTM 64 PLUS system (Micromed S.p.A., Mogliano Veneto, Italy). Each patient had 14 to 17 electrodes implanted, with 5 to 18 contacts per electrode. In one patient, radiofrequency thermocoagulation (RFTC) was performed at the end of the SEEG examination. Participation in the present study had no influence on the clinical planning or execution of the SEEG investigations, and did not influence the decision to perform RFTC.

### 2.3 SEEG analysis

Representative electrode samples for omics analyses were selected prior to explantation based on the visual inspection of SEEG signals, following current clinical practice (Isnard et al., 2018). Representative samples were chosen from three electrophysiologically distinct zones:

1. Epileptogenic zone (EZ), defined as the brain area considered necessary for resection to achieve seizure freedom, primarily based on the earliest onset and organization of ictal discharges during spontaneous habitual seizures. Additional information, including interictal epileptiform activity and stimulation-induced seizures, was considered.
2. Propagation zone (PZ), defined as brain regions showing the earliest propagation of ictal discharges outside the epileptogenic zone.
3. Least involved zone (LIZ), defined as areas without ictal and interictal epileptic activity, or areas involved only in the late spread of ictal discharges and lacking independent interictal epileptiform activity.

For each patient, up to four samples from each zone were included and allocated to transcriptomic and proteomic analyses (see table 2). The minimum sample length corresponded to three consecutive electrode contacts. Electrode contacts subjected to RFTC were not selected as samples.

**Table 2.**
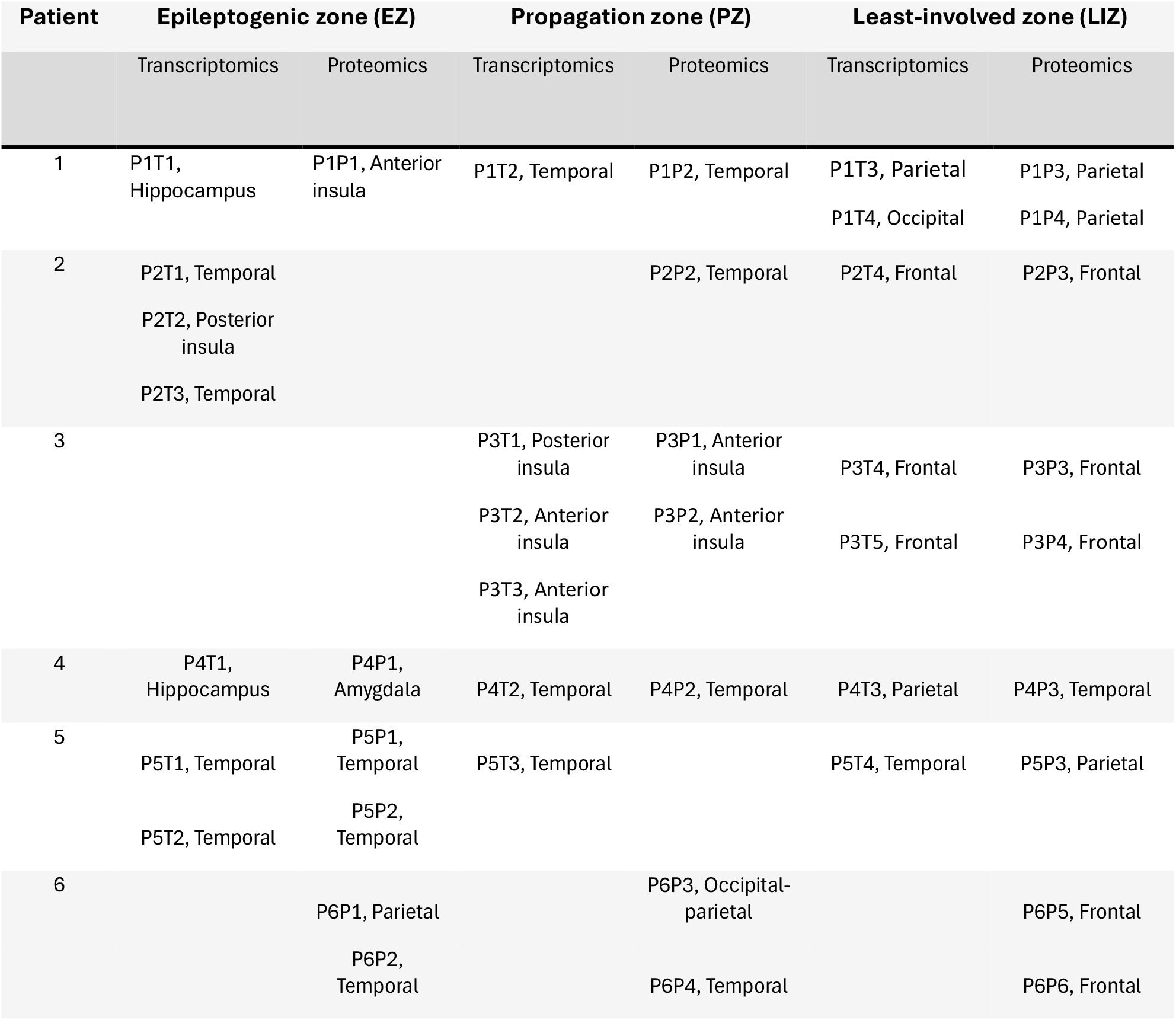
Electrophysiological and anatomical characteristics of samples used for transcriptomic and proteomic analyses. Sample codes and anatomical locations are reported. In this table, Temporal refers to the neocortical temporal cortex. Abbreviations: PnTm, patient n transcriptomics sample m; PnPm, patient n proteomics sample m.

| Patient | Epileptogenic zone (EZ) |  | Propagation zone (PZ) |  | Least-involved zone (LIZ) |  |
| --- | --- | --- | --- | --- | --- | --- |
|  | Transcriptomics | Proteomics | Transcriptomics | Proteomics | Transcriptomics | Proteomics |
| 1 | P1T1, Hippocampus | P1P1, Anterior insula | P1T2, Temporal | P1P2, Temporal | P1T3, Parietal<br>P1T4, Occipital | P1P3, Parietal<br>P1P4, Parietal |
| 2 | P2T1, Temporal<br>P2T2, Posterior insula<br>P2T3, Temporal |  |  | P2P2, Temporal | P2T4, Frontal | P2P3, Frontal |
| 3 |  |  | P3T1, Posterior insula<br>P3T2, Anterior insula<br>P3T3, Anterior insula | P3P1, Anterior insula<br>P3P2, Anterior insula | P3T4, Frontal<br>P3T5, Frontal | P3P3, Frontal<br>P3P4, Frontal |
| 4 | P4T1, Hippocampus | P4P1, Amygdala | P4T2, Temporal | P4P2, Temporal | P4T3, Parietal | P4P3, Temporal |
| 5 | P5T1, Temporal<br>P5T2, Temporal | P5P1, Temporal<br>P5P2, Temporal | P5T3, Temporal |  | P5T4, Temporal | P5P3, Parietal |
| 6 |  | P6P1, Parietal<br>P6P2, Temporal |  | P6P3, Occipital-parietal<br>P6P4, Temporal |  | P6P5, Frontal<br>P6P6, Frontal |

Prior to omic analyses, the final SEEG reports, usually available only after explantation, were reviewed, and sample classifications were adjusted when necessary. For patient 3, the epileptogenic zone could not be conclusively defined. Details of all analyzed SEEG electrode samples and their final classifications are provided in Table 2.

### 2.4 Sample collection

Following electrode removal, the preselected SEEG electrode contacts were cut and placed into individually labeled tubes (Figure 1). For transcriptomic analyses, electrodes from patients 1-3 were processed immediately after explantation and transported on ice to a nearby laboratory for sample preparation. Based on the initial experience, the protocol was optimized for patients 4 and 5 by briefly immersing the electrodes in cold phosphate-buffered saline (PBS) prior to cutting to remove residual blood and other surface contaminants.

**Figure 1.**
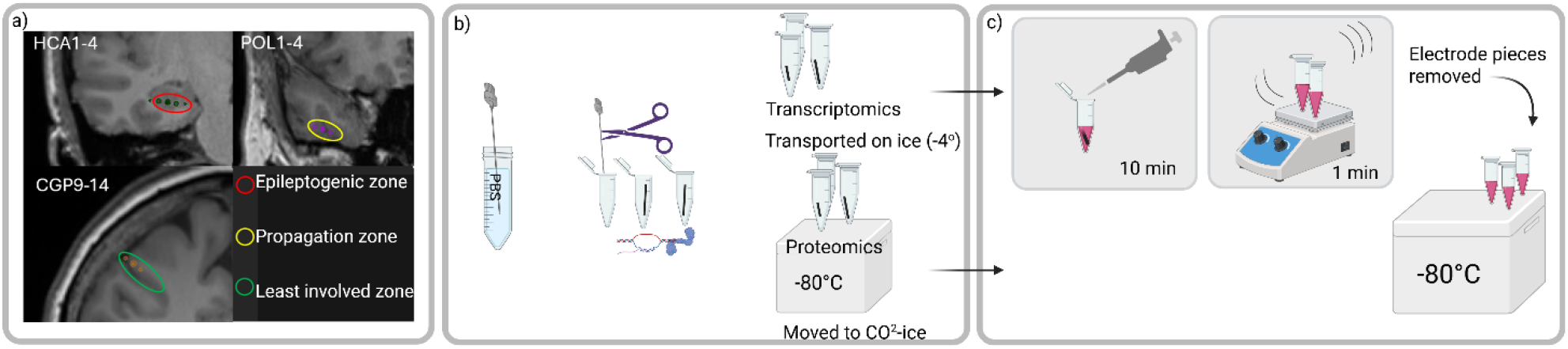
Overview of the final workflow for the collection and processing of SEEG-derived human brain tissue for transcriptomic and proteomic analyses. The figure illustrates SEEG analysis and sample selection (a), sample collection (b), and sample preparation (c). Procedures were performed at the Department of Surgery, New Children’s Hospital (pediatric patients), and the Department of Neurosurgery, Helsinki University Hospital, Helsinki, Finland (adult patients).

For proteomic analyses, electrodes from patients 1-3 were cut and placed into tubes containing cold PBS before being transported on ice to the laboratory. For patients 4-6, the optimized protocol involved briefly rinsing the electrodes in cold PBS, cutting the selected contacts into collection tubes in the operating room, immediately placing the tubes on dry ice, and subsequently storing them at −80°C until further processing (Figure 1).

To assess the presence of adherent brain tissue, cytological analysis was performed on a representative SEEG electroe from one patient. The electrode was vortexed in physiological saline, and the resulting cell suspension was fixed in ethanol. Cytospin preparations were generated and stained using the Papanicolaou method for microscopic evaluation.

### 2.5 Sample preparation on site for RNA isolation

Samples from patients 1-5 were prepared for cryopreservation in a laboratory located within the surgery department within one hour of electrode removal. Each sample was incubated in 700 µl of QIAzol Lysis Reagent (QIAGEN, Hilden, Germany) at room temperature for 10 minutes and subsequently vortexed briefly. After removal of the electrode segment, the lysates were flash-frozen on dry ice and stored at −80°C until further analyses. The protocol required only standard laboratory equipment, including a fume hood, vortex mixer, and dry ice, facilitating implementation in a clinical surgical ward.

### 2.6 Sample preparation on site for liquid chromatography–tandem mass spectrometry (LC-MS/MS) analysis

For patients 1-3, samples designated for proteomic analysis were processed in the laboratory of the surgery department. Following transport on ice, the samples were vortexed for 3–6 s and centrifuged at 400 × *g* for 15 min at 4°C to pellet the tissue material. The resulting pellets were then flash-frozen and stored at −80°C until further analysis.

For patients 4–6, an optimized workflow was implemented in which electrode samples were immediately placed on dry ice in the operating room following collection and subsequently stored at −80°C until LC–MS/MS analysis.

### 2.7 RNA isolation, preparation, and sequencing

RNA isolation, quality control, library preparation and next-generation sequencing (NGS) were performed by the HiPREP Core Facility at the FIMM Technology Centre and the Genomics NGS Sequencing unit at the University of Helsinki. Detailed protocols for RNA isolation, RNA elution, RNA integrity assessment, library preparation and sequencing are provided in Appendix S1. Downstream analyses for transcriptomics are summarized in supplementary files S3.

### 2.8 LC–MS/MS and analysis

For samples from patients 1-3, 500 µl of 8 M urea (Fisher, cat. no. 424585000) in 100 mM AMBIC (ammonium bicarbonate (NH4HCO3, #A6141, Sigma Aldrich)) was added into the solution and for samples from patients 4-6, 300 µl of the same buffer was added directly to the electrode segments. From this stage onward, all samples were processed using an identical workflow. Detailed sample preparation procedures and details of subsequent analyses are provided in Appendix S2. Downstream analyses for proteomics are summarized in supplementary files S3.

### 2.9 Ethical aspects

The research plan was approved by the Ethics Commission of Helsinki University Hospital (HUS/7438/2023), and the study was conducted in accordance with the Declaration of Helsinki. During the SEEG investigation, patients were invited to participate in the study, and informed consent was obtained from the patients or their legal guardians prior to sample collection.

## 3 Results

### 3.1 Brain tissue attached to SEEG electrodes as a source for transcriptomic and proteomic analyses

Transcriptomic and proteomic analyses were successfully completed for all collected samples, demonstrating that SEEG-derived tissue can support comprehensive molecular profiling. Optimization of the tissue collection protocol during the study had no substantial effect on the number of detected transcripts or proteins per sample. The overall performance of the workflow remained robust across protocol iterations.

Cytological evaluation demonstrated presence of a small number of degenerated cells, likely representing neuronal and/or glial cells, together with extracellular matrix material and red blood cells.

All samples yielded sufficient RNA for sequencing, with RNA concentrations ranging from 2.18 to 30.69 ng/µL. RNA quality was generally high, with RNA Integrity Number (RIN) values exceeding 7 in all samples except P4T1, which had a RIN value of 5.1.

In the LC-MS/MS proteomics analysis, a total of 5,717 unique proteins were identified across all samples. Following filtering to retain proteins detected in at least 30% of samples, 4,592 proteins remained for downstream analyses. The number of detected proteins was consistent across samples and protocol iterations, indicating stable technical performance of the proteomics workflow (figure S4).

To assess the major sources of variability within the datasets, principal component analysis (PCA) was performed. In the transcriptomic dataset, the first principal component (PC1) primarily separated samples according to anatomical origin (Figure 2A). Neocortical temporal lobe samples formed a relatively tight cluster, whereas anterior and posterior insula samples appeared more distinct, with partial clustering observed between insular subregions. The second principal component (PC2) reflected patient-specific variation.

**Figure 2.**
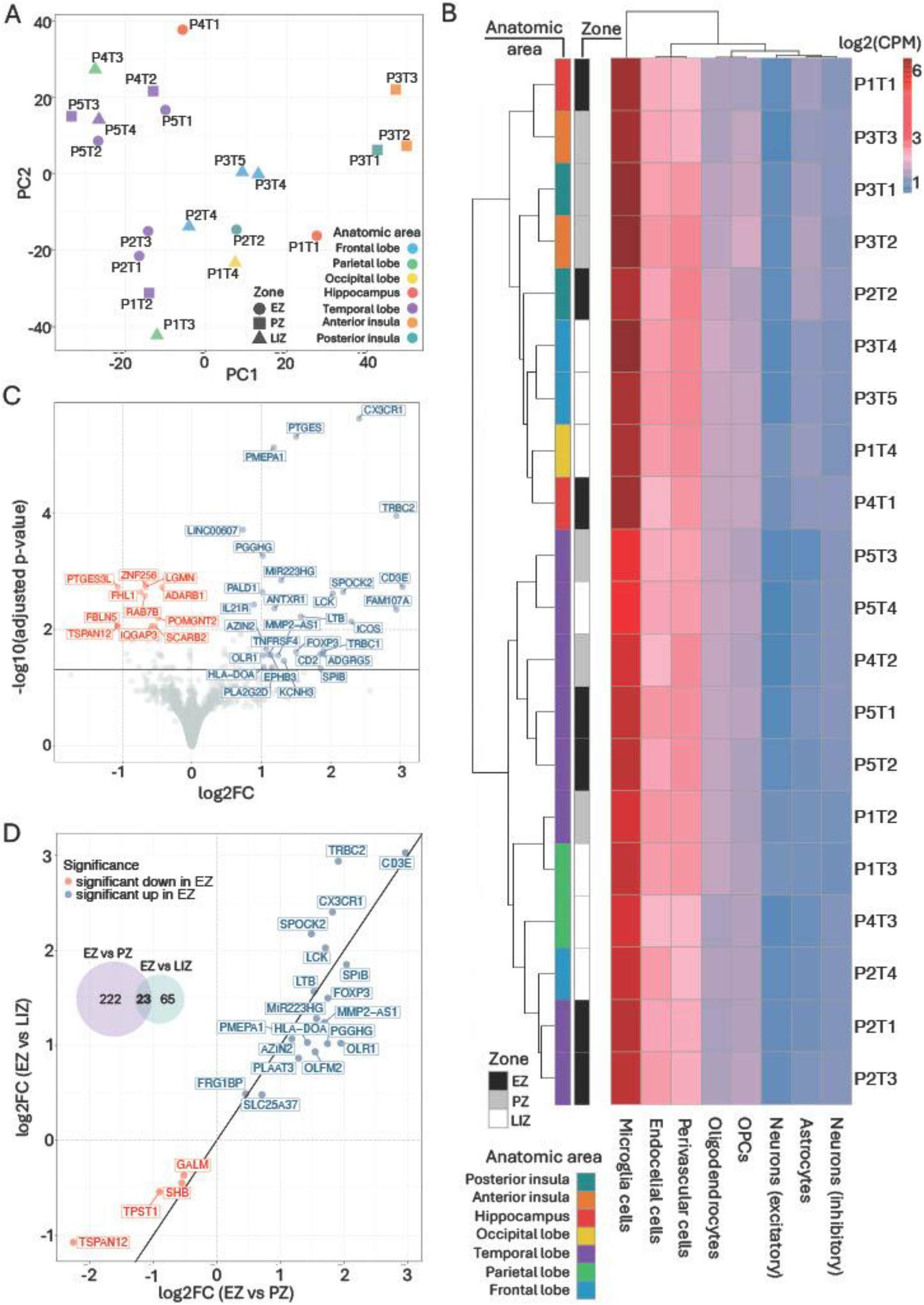
(A) Principal component analysis of transcriptomic samples. Points are colored by anatomical area and shaped according to the electrophysiological zone. (B) Hierarchically clustered heatmap showing log2 fold change of counts per million (log2FC, CPM) of cell type– specific marker genes across transcriptomic samples. Sample color annotations indicate anatomical area and zone. (C) Volcano plot of differential gene expression between EZ and LIZ samples. Genes with higher absolute log2FC or lower adjusted p-values are annotated. (D) Scatter plot comparing log2FC values from the EZ versus PZ and EZ versus LIZ differential expression analyses. In this figure, Temporal refers to neocortical temporal cortex.

A similar pattern was observed in the proteomic dataset. While no clear separation based on anatomical region was evident, PC2 again captured a pronounced patient-specific effect (Figure S5), which emphasized the importance of accounting for individual variability in subsequent analyses.

### 3.2 Anatomical region–driven clustering of cell-type–specific transcriptional signatures

To characterize major brain cell-type signatures within the SEEG transcriptomic dataset, we leveraged cell-type annotations derived from the HPA (Figure 2B). Consistent with the PCA results, the overall clustering pattern was driven primarily by anatomical origin rather than epileptic zone classification. Two major sample clusters were identified: one predominantly composed of neocortical temporal and parietal lobe samples, and a second cluster enriched for samples from other brain regions, including frontal cortex, insula, and hippocampus. This latter group showed a relative enrichment of microglia-associated transcriptional signatures.

### 3.3 EZ displays the strongest transcriptional divergence among sampled regions

Differential expression analysis revealed substantial transcriptional differences across zones, with the EZ showing the most distinct molecular profile. Comparison of the EZ and LIZ identified 88 significantly differentially expressed genes, including 50 upregulated and 38 downregulated genes in the EZ (Table S1). Among the most strongly upregulated genes were *CX3CR1, TRBC2* and *CD3E*, while *PTGES3L, TSPAN12*, and *FBLN5* showed the strongest reductions in expression (Figure 2C). Several of the top differentially expressed genes, including *CX3CR1, PTGES, PMEPA1*, and *TRBC2*, were associated with immune and inflammatory processes. In contrast, genes enriched in the LIZ were fewer in number and generally exhibited smaller effect sizes.

An even stronger transcriptional divergence was observed between the EZ and PZ, with 245 significantly differentially expressed genes identified, including 116 upregulated and 129 downregulated genes in the EZ (Table S1, Figure S6). Genes most strongly enriched in the EZ included *GFAP, CX3CR1, CD3E, CD69*, and several HLA class II–related genes (*HLA-DQA2, HLA-DQB2, and HLA-DOA*), consistent with increased glial, immune, and inflammatory activity. In contrast, relatively few genes were strongly enriched in the PZ, with *TSPAN12, ADAMTS15, LRP5*, and *GPRC5B* among the most prominent.

We next examined genes that were significant in both the EZ versus PZ and EZ versus LIZ comparisons (Figure 2D). All of the shared genes exhibited consistent direction and magnitude of differential expression across both comparisons. Particularly, TRBC2, CD3E, and CX3CR1 were among the most consistently upregulated genes, whereas *TSPAN12, TPST1, GALM* and *SHB* showed downregulation in the EZ.

### 3.4 Immune and antigen-presentation pathways dominate the epileptogenic zone transcriptome

Pathway enrichment analysis revealed a strong immune-associated signature within the EZ (Figure 3A, Table S2). The most significantly enriched biological processes pathways included T-cell activation and regulation, lymphocyte differentiation, cytokine-mediated signaling, and other pathways associated with adaptive immune responses. Enriched cellular component pathways were predominantly related to the extracellular matrix, plasma membrane, cell surface, and cell periphery, which can suggest structural remodeling and altered cell–cell communication within epileptic tissue. Molecular function pathways further supported these findings, with significant enrichment of receptor activity and signaling-related functions.

**Figure 3.**
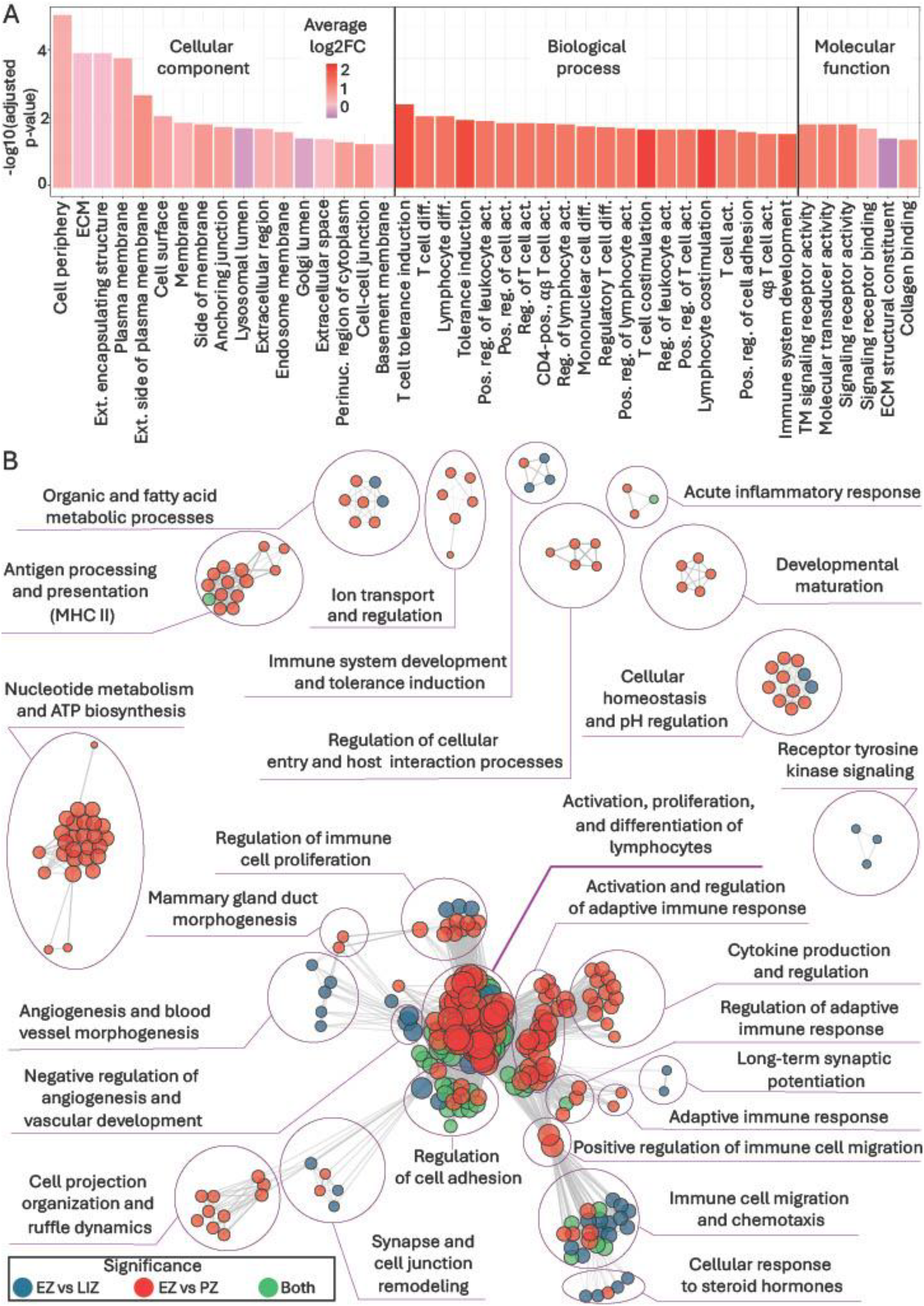
(A) ORA enrichment analysis of differentially expressed genes comparison between EZ and LIZ. Significantly enriched pathways are grouped into the GO categories. For biological processes, top 20 significant pathways were selected based on highest absolute average expression of related genes. Abbreviations: ECM, extracellular matrix; ext., external; perinuc., perinuclear; diff., differentiation; pos., positive; reg. regulation; act., activation; (B) Enrichment map of significant pathways annotated by GO as biological processes. Nodes represent enriched GO pathways and edges indicate shared genes between the nodes.

An even stronger enrichment pattern was observed in the comparison between EZ and PZ (Figure S7, Table S2). Majority of significant pathways were mostly centered on immune and inflammatory functions. Within the cellular component ontology, highly enriched pathways included MHC class II protein complex, MHC protein complex, and cell-surface compartments, which can indicate enhanced antigen presentation capacity within the epileptogenic region. Similarly, biological process pathways were enriched for T-cell activation, lymphocyte differentiation, monocyte chemotaxis, and interferon-mediated signaling. Molecular function associated pathways identified MHC class II receptor activity, antigen binding, and immune receptor activity among the most significantly enriched.

Consistent with the GO findings, KEGG pathway analysis demonstrated a strong enrichment of immune- and inflammation-related pathways in the EZ (figure S8). Furthermore, the EZ versus PZ comparison revealed a relative suppression of multiple metabolic pathways, indicating that immune activation in epileptogenic tissue is accompanied by substantial metabolic remodeling (figure S9).

To determine whether the pathway-driven alterations observed in the EZ represented shared or comparison-specific biological processes, semantic similarity network analysis was performed across significantly enriched GO pathways identified in the EZ versus LIZ and EZ versus PZ comparisons (Figure 3 B). This analysis revealed extensive overlap between the two comparisons. Analysis of cellular component GO ontology pathways further supported these observations (Figure S10). Shared enrichments were largely associated with membrane- and lysosome-related compartments, indicating common alterations in intracellular trafficking and membrane organization within the EZ. Molecular function associated pathway analysis revealed a similar pattern too (Figure S11). Functional alterations shared between comparisons were relatively few, whereas the EZ versus PZ comparison showed extensive enrichment of immune-related receptor and binding activities associated with antigen recognition, cytokine signaling, and adaptive immune responses.

### 3.5 Transcriptomic and proteomic changes exhibit directional concordance across comparisons between electrophysiological zones

In contrast to the transcriptomic analysis, which identified a substantial number of differentially expressed genes across comparisons, no proteins reached statistical significance in the proteomic dataset following multiple testing correction (Table S3). Given this limitation, the proteomic data were not interpreted in isolation and instead we focused on assessing the consistency of directionality between transcriptomic and proteomic changes by comparing log2FC for genes that were significantly differentially expressed at the transcript level and mapped to corresponding proteins. This comparison, for both the EZ versus LIZ and EZ versus PZ contrasts, demonstrated strong directional concordance between the two data modalities (Figures S12, S13).

## 4 Discussion

This study describes a practical and robust method for performing omic studies on tissue adhered to SEEG electrodes. The sample collection and preprocessing protocol was successfully performed by clinical investigators in a standard surgical setting. These samples generated high-quality transcriptomic and proteomic datasets suitable for downstream analyses.

Previous studies investigating epileptogenic tissue in rodent models and human surgical or postmortem specimens also employed omic methodologies. While resected human tissue obtained during epilepsy surgery provides valuable knowledge into the local epileptogenic zone, it does not capture molecular alterations in the broader epileptic network, where secondary epileptogenesis may take place. Furthermore, rodent models are unable to fully capture the complexity of epileptogenic processes in the human brain (Becker, 2018). Comparisons with postmortem human brain tissue are similarly challenging due to concerns regarding RNA degradation and reduced transcript stability (Dachet et al., 2021). In contrast, the use of SEEG electrodes for omic analyses enables the investigation of multiple regions within the epileptic network, as well as areas outside it, allowing patients to serve as their own controls.

Recently, (Dwivedi et al., 2025) demonstrated the feasibility of RNA sequencing and epigenomic profiling using tissue collected from SEEG electrodes through an alternative sampling approach. They immediately snap-froze electrodes following removal, obtaining RNA with RIN values ranging from 1.2 to 8.4 (mean 5.9). In the present study, samples were immersed in QIAzol within 10–20 minutes after electrode removal. RNA isolated using our protocol exhibited slightly higher quality, with RIN values ranging from 5.1 to 9.7 (mean 8.8). However, differences in downstream laboratory procedures may also have contributed to the observed variation in RNA quality between studies.

Due to the small and heterogenous sample of patients, we did not perform an in-depth analysis of individual differentially expressed transcripts and proteins, nor did we systematically compare our molecular findings with those reported by Dwivedi et al., (2025). Nevertheless, the datasets generated revealed several important observations that warrant further investigation in larger, more homogeneous patient cohorts.

In the transcriptomic analyses, differences between the EZ and the outside area (PZ and LIZ) suggest that the EZ is characterized by a strong neuroinflammatory response, and a reduced metabolic capacity and mitochondrial function. These preliminary findings are consistent with the findings supporting the involvement of immune-mediated mechanisms in drug-resistant epilepsy (Shi et al., 2025; Solanki & Jha, 2025) and with reports describing altered cellular homeostasis and metabolic adaptation within the epileptic focus in chronic epilepsy (Liotta et al., 2024). In contrast, the comparison between PZ and LIZ did not reveal any significantly differentially expressed genes, possibly reflecting both the limited sample size and the heterogeneity of the epilepsy etiologies included in this pilot study. At the same time, the observed differences in inflammatory signaling between the EZ and the other regions suggest that these responses are unlikely to be solely attributable to SEEG implantation and may instead reflect biological processes specific to the epileptogenic zone.

In contrast to the transcriptomic analysis, which identified a substantial number of differentially expressed genes in the EZ in comparison to other areas, no proteins remained statistically significant in the proteomic dataset following correction for multiple testing. This discrepancy is likely attributable to the limited sample size and the consequently reduced statistical power of the proteomic analysis, as well as the inherently higher variability and lower dynamic range typical of mass spectrometry–based protein quantification. However, the proteomic data were of high quality, and a substantial number of proteins were successfully quantified across samples, indicating considerable potential for detecting biologically meaningful protein-level alterations in future studies with larger cohorts.

Several limitations of this proof-of-concept study should be acknowledged. The number of patients included was small, and the underlying etiology of epilepsy varied across patients. Additional potential confounding factors include age, sex, the anatomical brain regions covered by SEEG investigations, and the extent and type of epileptic activity preceding sample collection. Furthermore, the sample collection method itself may introduce variability, as electrode contacts traverse multiple brain regions during explantation. To minimize the potential impact of blood contamination and non-adherent debris, we introduced a flushing step prior to processing, thereby enriching the samples for tissue attached directly to the electrode surface. The comparison of flushed and non-flushed samples revealed no significant differences in the number of unique molecular identifiers detected, suggesting that this procedure does not adversely affect sample yield.

## Conclusions

Overall, our findings demonstrate that SEEG electrodes represent a valuable source of biological material for molecular characterization of chronic epilepsy in the living human brain. Future studies involving larger patient cohorts will facilitate more detailed comparisons across epilepsy etiologies and electrophysiological zones, while direct comparison with surgical resection specimens will provide further validation of this method. This method has the potential to facilitate the identification of both shared and distinct molecular mechanisms underlying epileptogenesis across patients with different epilepsy etiologies.

## Acknowledgments

The project was supported by Päivikki and Sakari Sohlberg Foundation, the Foundation for Pediatric Research, and the Research Council of Finland (decision numbers 369453 and 369391). Personal grants have been awarded for Ninja Ehrnrooth from Arvo and Lea Ylppö foundation and the Finnish Epilepsy research foundation. The authors would like to thank HiPREP Core at FIMM Technology Centre supported by HiLIFE, Biocenter Finland for the RNA isolation and quality control and FIMM Genomics NGS Sequencing, Institute for Molecular Medicine Finland, FIMM for the sample sequencing and the Molecular Systems Biology Research Group & Proteomics unit HiLIFE; Institute of Biotechnology for the LC–MS/MS and proteomic analysis. We also want to thank the Clinical Trials Unit in HUS and Dr. Jaakko Klockars for support.

