## Supplemental File for "A novel method for obtaining epileptic brain tissue for omic analyses using electrodes from a clinical stereoelectroencephalography study"

### Supplement

#### S1, RNA isolation, library preparation and sequencing

Total RNA was isolated with the miRNeasy micro kit (QIAGEN, Hilden, Germany) according to the manufacturer's instructions including the DNAse I treatment in the QIACube instrument.

RNA integrity was assessed using the High Sensitivity RNA ScreenTape (Agilent Technologies) in the Agilent 4200 TapeStation System (Agilent Technologies, Santa Clara, CA, USA). The concentration was measured by QuantiFluor® RNA System (Promega, Madison, WI, USA).

cDNA was synthesized from 10 ng of total RNA using the SMART-Seq® mRNA Kit (Takara Bio, Mountain View, CA, USA) according to the manufacturer's instructions. Sequencing libraries were subsequently prepared using the Nextera XT DNA Library Preparation Kit (Illumina, San Diego, CA, USA) following the Illumina Nextera XT Reference Guide. Unique dual indexing (UDI) was used for library preparation.

Library quality check was performed using Agilent Fragment Analyzer HS NGS Fragment kit (Agilent, Santa Clara, CA, USA). Libraries were pooled based on the concentrations determined during quality control and quantified using the Collibri™ Library Quantification Kit (Thermo Fisher Scientific, Waltham, MA, USA).

Sequencing was performed on an Illumina NovaSeq 6000 platform using an S1 flow cell and a 200-cycle sequencing kit (Illumina, San Diego, CA, USA). Paired-end sequencing was carried out with a read length of 2 × 101 bp.

#### S2, Laboratory steps for sample preparation and methods for LC–MS/MS analysis

Samples were incubated on ice for 6 hours with two 10-minute sonications and intermittent vortexing. Lysed samples were reduced with 5 mM Tris(2-carboxyethyl) phosphine hydrochloride (Thermo Scientific, cat. #20490) for 30 minutes in 37 °C and alkylated with 10 mM iodoacetamide (Acros Organics, cat. #122271000) for 30 minutes at room temperature, and the urea concentration was diluted to 1 M with 100 mM AMBIC. Reduced and alkylated peptides were trypsin-digested at 37°C for 16 hours using Sequencing Grade Modified Trypsin (V5113, Promega). After digestion, samples were acidified with 10% trifluoroacetic acid (TFA, #85049.051, VWR) and desalted with BioPureSPN PROTO 300 C18 Mini columns (HUM S18V, Higgins analytical, USA) according to manufacturer's instructions. Then desalted samples were dried in a centrifuge concentrator (Concentrator Plus, Eppendorf). The dried peptides were reconstituted in 30 µl buffer A (0.1% (vol/vol) TFA, 1% (vol/vol) acetonitrile (83640.320, VWR) in HPLC grade water (#10777404, Fisher Scientific)). Samples were further diluted 10+10 µl with HPLC water containing 0.1 vol/vol% formic acid. The manufacturer's instructions were followed to load into Evotips (EV2003, EvoSep, Denmark).

Proteomic analyses were performed with dia-PASEF using the Evosep One (Evosep, Denmark) coupled to a Bruker timsTOF Pro2 (Bruker Daltonics) (Meier et al., 2018, 2020) via a CaptiveSpray nano-electrospray ion source (Bruker Daltonics). A 15 cm × 75 µm column with 1.9 µm C18 beads (EV1112, Evosep) was used for peptide separation with the 40 samples per day Whisper Zoom method (32.5 min gradient time). Mobile phases A and B were 0.1% formic acid in water and 0.1% formic acid in acetonitrile, respectively. The MS analysis was performed in the positive-ion mode with dia-PASEF (Meier, Brunner et al., 2020) method with sample optimized data independent analysis (dia) scan parameters. We performed DDA in PASEF mode from a pooled sample to be able to adjust dia-PASEF parameters optimally. To perform sample-specific dia-PASEF parameter adjustment, the default dia-short-gradient acquisition methods were adjusted based on the sample specific DDA-PASEF run with the software “tims Control” (Bruker Daltonics). The ion mobility windows were set to best match the ion cloud density from the sample type-specific DDA-runs. The following parameters were used in dia-PASEF runs: TIMS settings 1/KO 0.85-1.30, Ramp time 100 ms, Accumulation time 100 ms, Duty cycle 100%, and dia-PASEF settings cycle time estimate 1.06 s, MS1 ramps 1, MS/MS ramps 9, MS/MS windows 24, mass range 430-1018.5 Da, Mobility range 0.85-1.22 1/K0. For collision energy at 0.60 1/KO 20 eV and at 1.60 1/KO 59 eV were used.

To analyze diaPASEF data, the raw data (.d) were processed with DIA-NN v1.9.2 (Demichev et al., 2020; Demichev et al., 2022) utilizing spectral library generated from the UniProt human proteome (UP000005640, downloaded 12.08.2024 as a FASTA file, 20410 proteins). During library generation, the following settings were used with fixed modifications: carbamidomethyl (C); variable modifications: acetyl (protein N-term), oxidation (M); enzyme:Trypsin/P; maximum missed cleavages: 1; mass accuracy fixed to 1.5e-05 (MS2) and 1.5e-05 (MS1); Fragment m/z set to 100-1700; peptide length set to 7-30; precursor m/z set to 300-1800; Precursor changes set to 2-4; protein inference not performed. All other settings were left to default. After DIA-NN, ambiguous protein groups were filtered out of the data, leaving only proteins quantified with unique peptides. Furthermore, proteins identified in less than 30% of the sample set were also filtered out. Samples were grouped into zone-based groups, and missing values were imputed using the QRILC method.Raw.d-files were processed with library-free search using DIA-NN v1.9.2 (Demichev et al., 2020). After DIA-NN, ambiguous protein groups were filtered out of the data, leaving only proteins quantified with unique peptides. Furthermore, proteins identified in less than 30% of the sample set were also filtered out. Samples were grouped into zone-based groups, and missing values were imputed using the QRILC method.

#### S3, Computational analysis

To characterize major brain cell-type signatures within the transcriptomic dataset, publicly available brain single-nucleus RNA sequencing annotations derived from the Human Protein Atlas (HPA) (Sjöstedt E et al., 2020) were used to define canonical cell-type clusters, which were subsequently aggregated into eight major cellular categories: astrocytes, oligodendrocytes, oligodendrocyte precursor cells (OPCs), microglia, excitatory neurons, inhibitory neurons, endothelial cells, and perivascular cells. For each category, marker genes were identified based on cluster-level expression profiles. Genes with high mean expression in the target cell type (nCPM > 100) and belonging to the top 1% of specificity scores relative to other cell types were selected. The top 20 most representative marker genes per cell type were retained for downstream analyses. These marker gene sets were then mapped onto the RNA-sequencing dataset. For each sample, a cell-type–specific expression score was computed by averaging the expression of the corresponding marker genes. These aggregated scores were used to generate a cell-type–resolved expression matrix across samples. The resulting matrix was visualized using hierarchical clustering heatmap to compare relative enrichment patterns across patients, sample zones and anatomical regions.

Differential gene expression analysis of the transcriptomic data was performed using the DESeq2 framework (Love et al., 2014). Prior to analysis, lowly expressed genes were filtered out by retaining only genes with at least 10 raw counts in a minimum of two samples. Samples were annotated according to the electrophysiological zone classification and grouped into three regions: EZ, PZ, and LIZ. To assess transcriptional differences between regions, pairwise comparisons were conducted using a generalized linear model implemented in DESeq2, with a design formula incorporating patient identity. The final model was fitted using negative binomial generalized linear modelling, and statistical testing for differential expression was performed using Wald test. P-values were adjusted for multiple testing using the Benjamini–Hochberg (BH) method, and genes with an adjusted p-value < 0.05 were considered significantly differentially expressed.

To facilitate biological interpretation of the differential expression results, over-representation analysis (ORA) was performed separately for each pairwise comparison of zones. Significantly enriched Gene Ontology (Thomas et al., 2022) pathways were identified and analyzed across the three GO ontology categories: Biological Process (BP), Molecular Function (MF), and Cellular Component (CC). In addition, the Kyoto Encyclopedia of Genes and Genomes (KEGG) collection of metabolic and signaling pathways was also used for pathway enrichment analysis (Kanehisa & Goto, 2000). P-values were adjusted for multiple testing using the BH method, and pathways with an adjusted p-value < 0.05 were considered significantly enriched.

To reduce redundancy among enriched pathways and identify functionally related biological groups, semantic similarity between GO pathways was calculated using the rrvgo package in R (Sayols, n.d.) .Pairwise semantic similarity scores were used to construct networks in which nodes represented enriched GO pathways and edges represented semantic relationships between them. Network analysis was subsequently applied to identify clusters of related GO pathways, and small clusters containing fewer than three pathways were excluded to improve interpretability. The resulting semantic similarity networks were visualized using force-directed graph layouts.

Differential protein intensity analysis was performed on log-transformed protein intensity data using the limma R package linear modeling framework. A linear model was fitted to compare zones of interest, while accounting for patient-specific effects as a factor to reduce inter-individual variability. Empirical Bayes moderation was applied to improve variance estimation across proteins, and differential intensity was assessed using moderated t-statistics (Gordon K Smyth, 2004). Multiple testing correction was performed using the BH method, and proteins with an adjusted p-value < 0.05 were considered significantly differentially expressed.

#### Supplemental figures

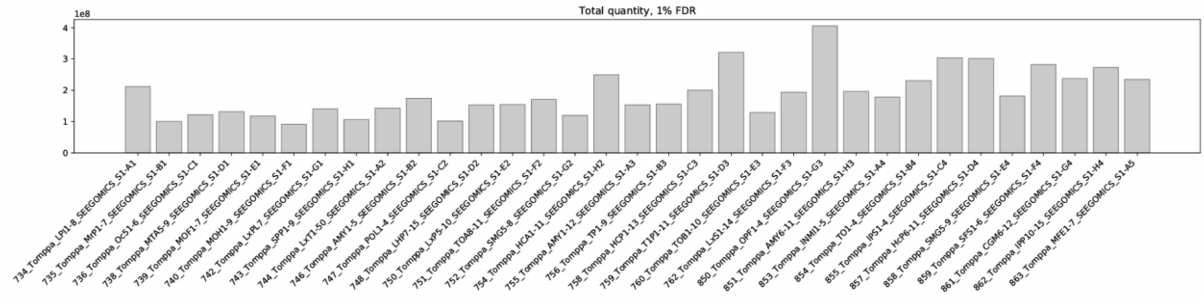


Figure S4 Total quantity of MS2 based precursors at 1% FDR.

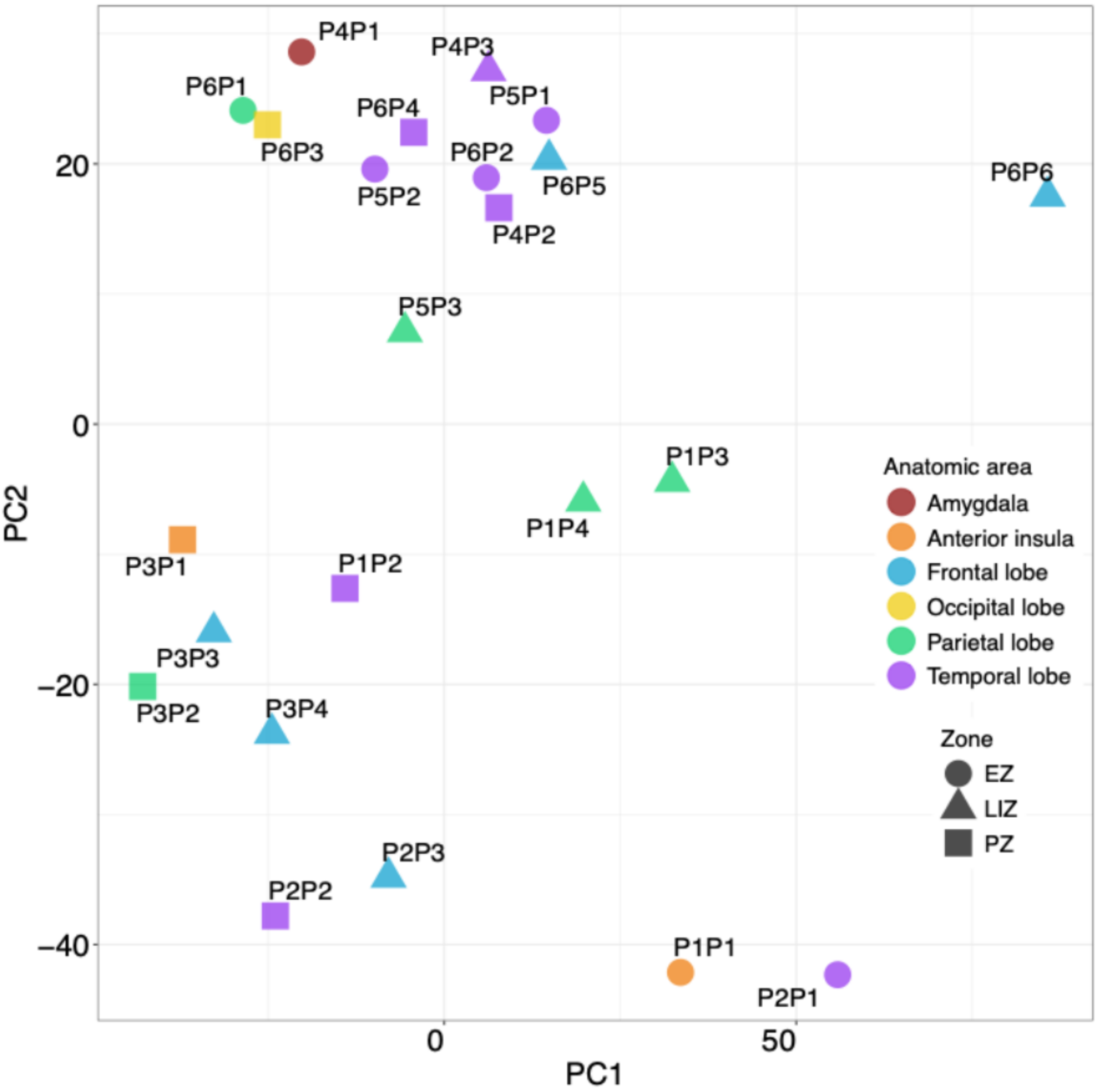


Figure S5 Principal component analysis of proteomic samples. Points are colored by anatomical area and shaped according to zone.

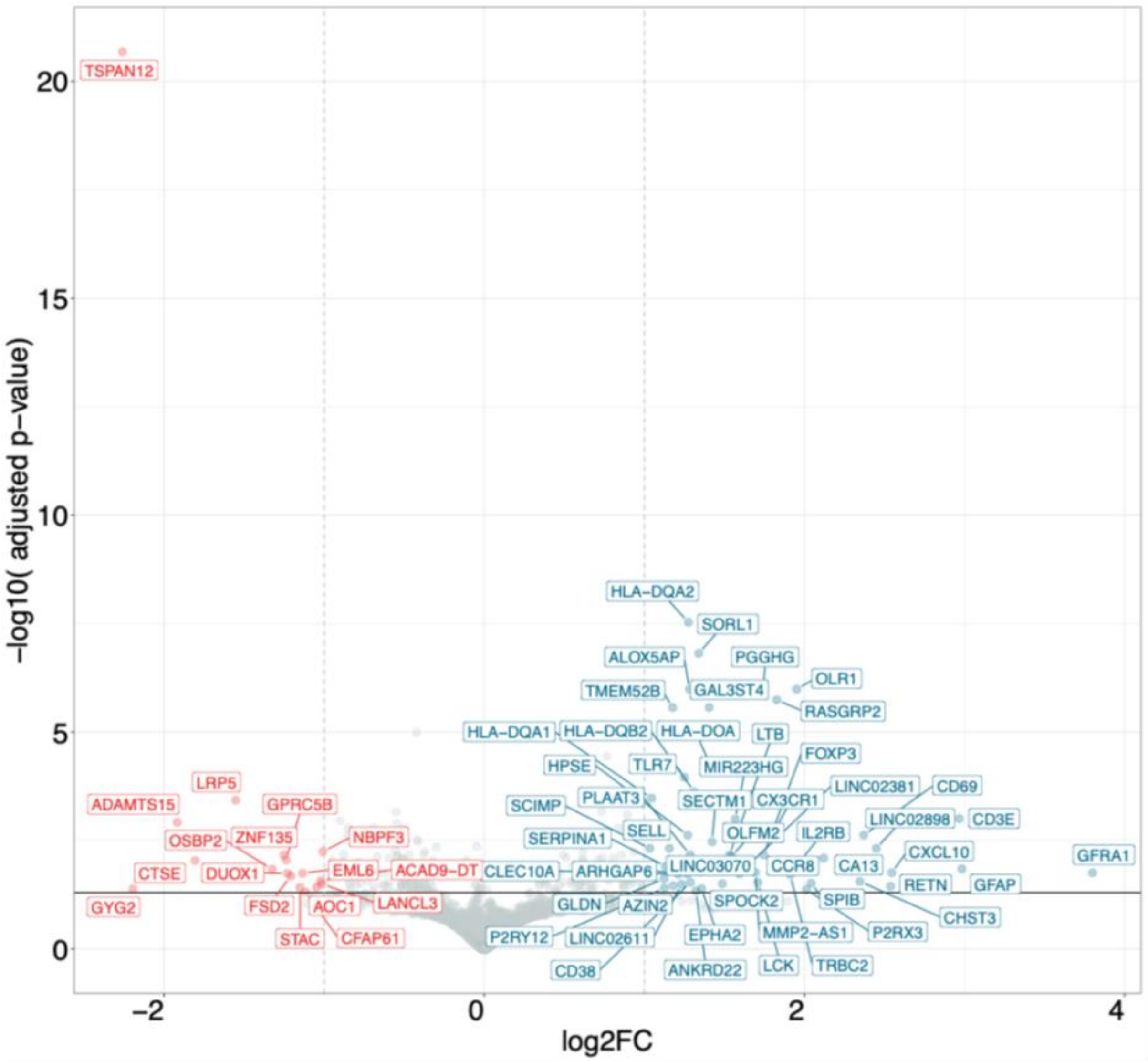


Figure S6 Volcano plot of differential gene expression between EZ and PZ samples. Significant genes with higher log2FC (absolute log2FC >1) are annotated.

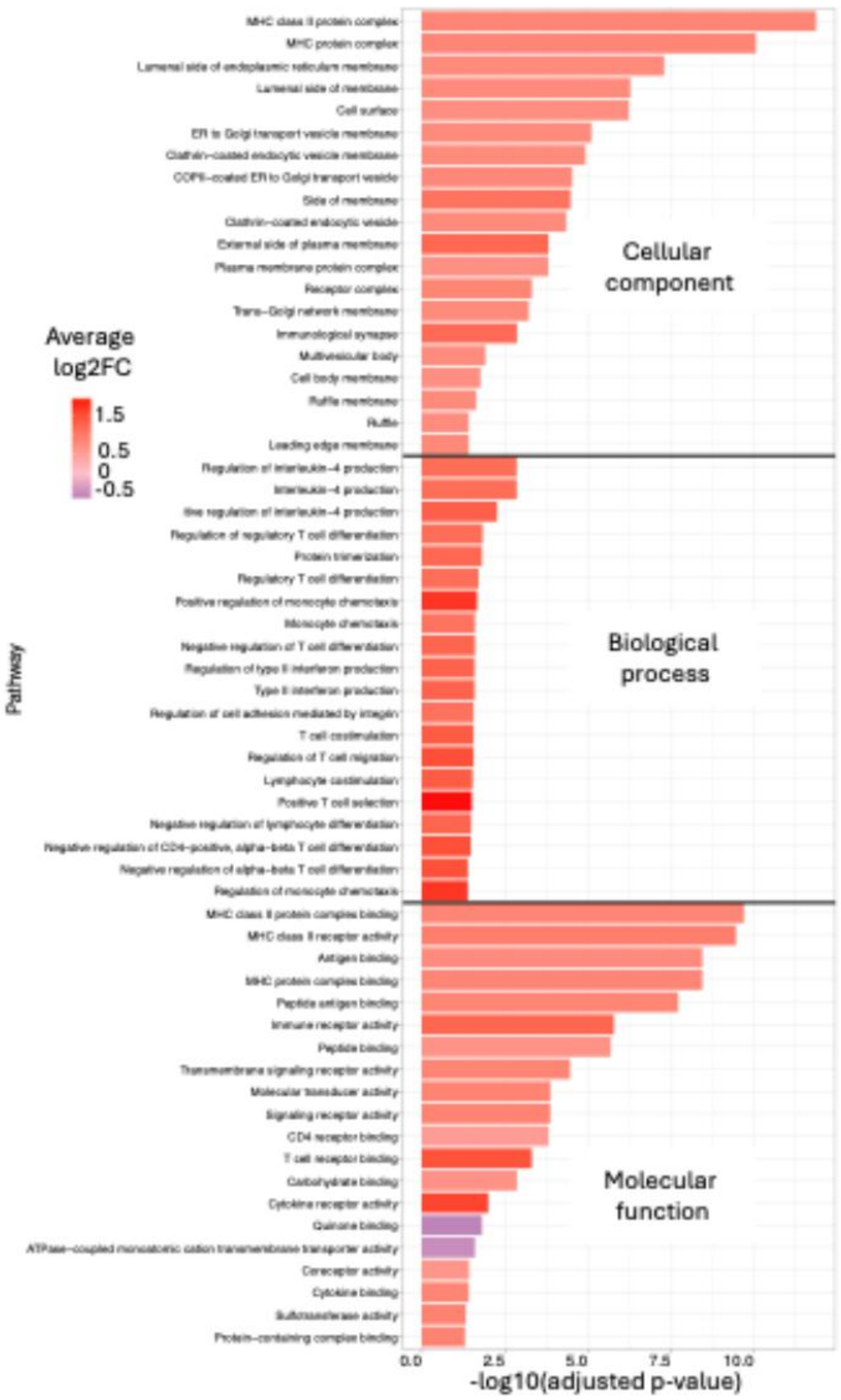


Figure S7 ORA enrichment analysis of differentially expressed genes in EZ versus PZ comparison. Significantly enriched pathways are grouped into the categories: cellular components, biological processes, and molecular functions. Top 20 significant pathways were selected per category based on highest absolute average expression of related significant genes.

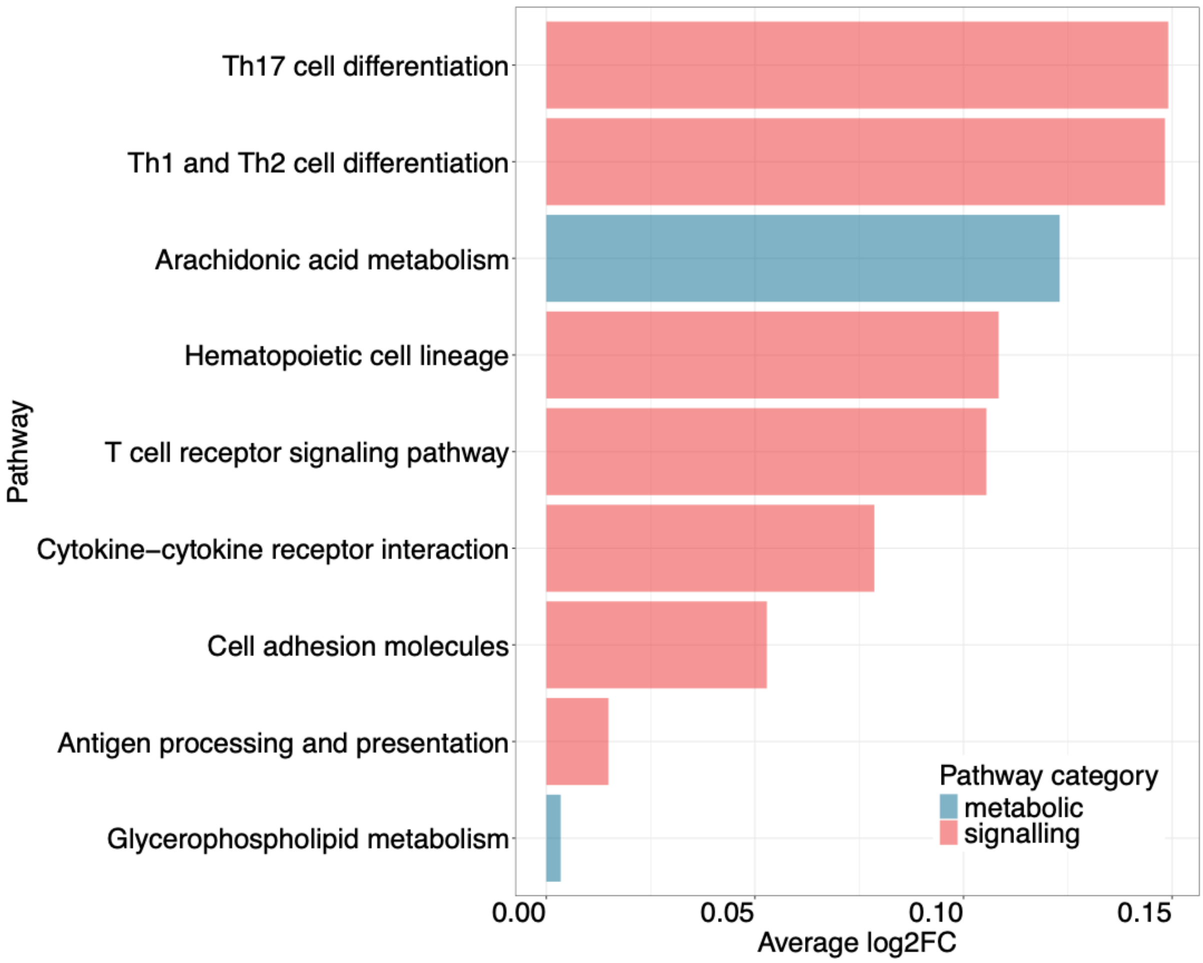

Figure S8 ORA enrichment analysis of differentially expressed genes in EZ versus LIZ comparison using KEGG pathway collection. Pathways are coloured based on categories.

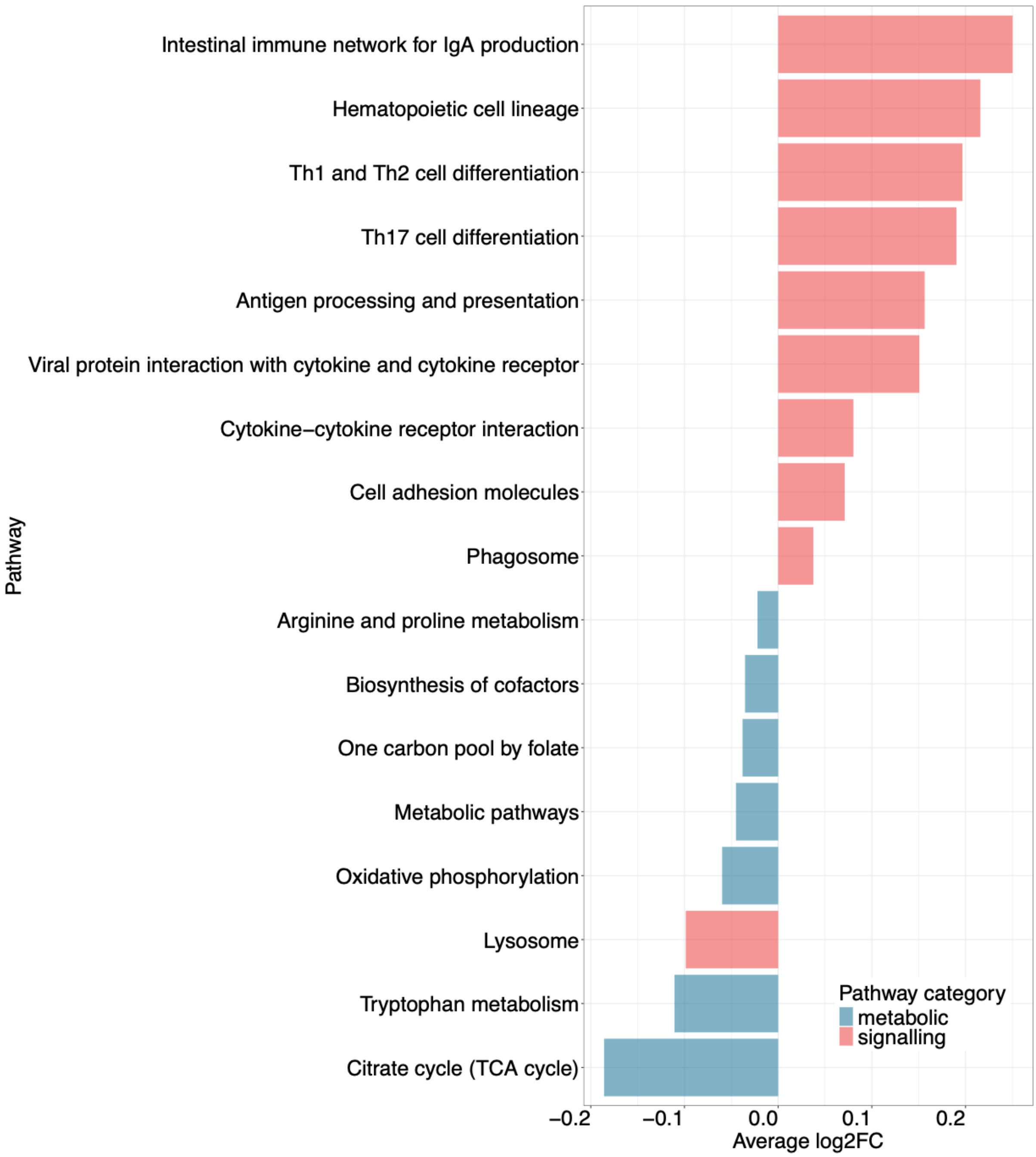


Figure S9 ORA enrichment analysis of differentially expressed genes in EZ versus PZ comparison using KEGG pathway collection. Pathways are coloured based on categories.

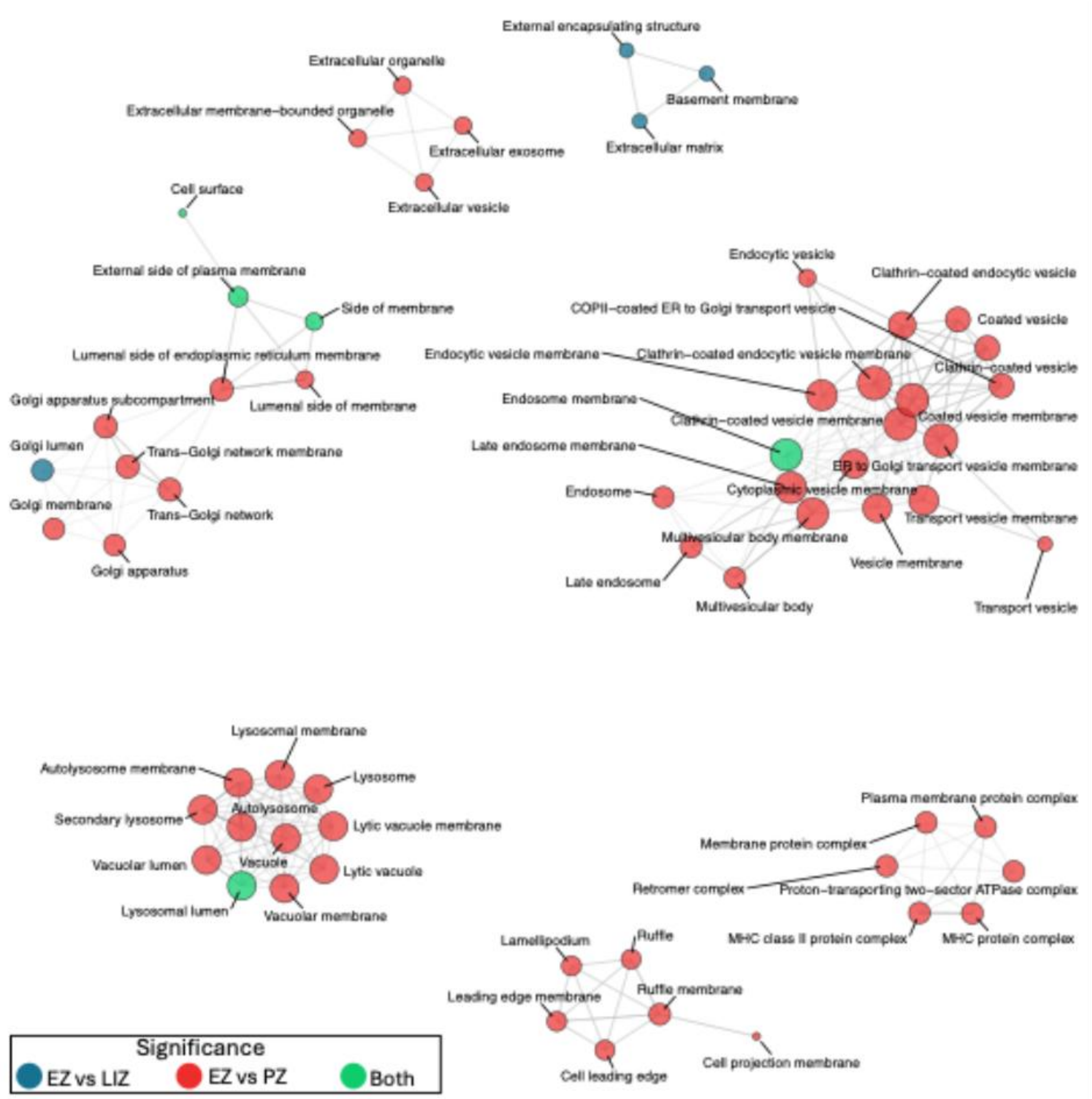


Figure S10 Enrichment map of significant pathways annotated by GO as cellular components. Nodes represent enriched GO pathways and edges indicate shared genes between the nodes.

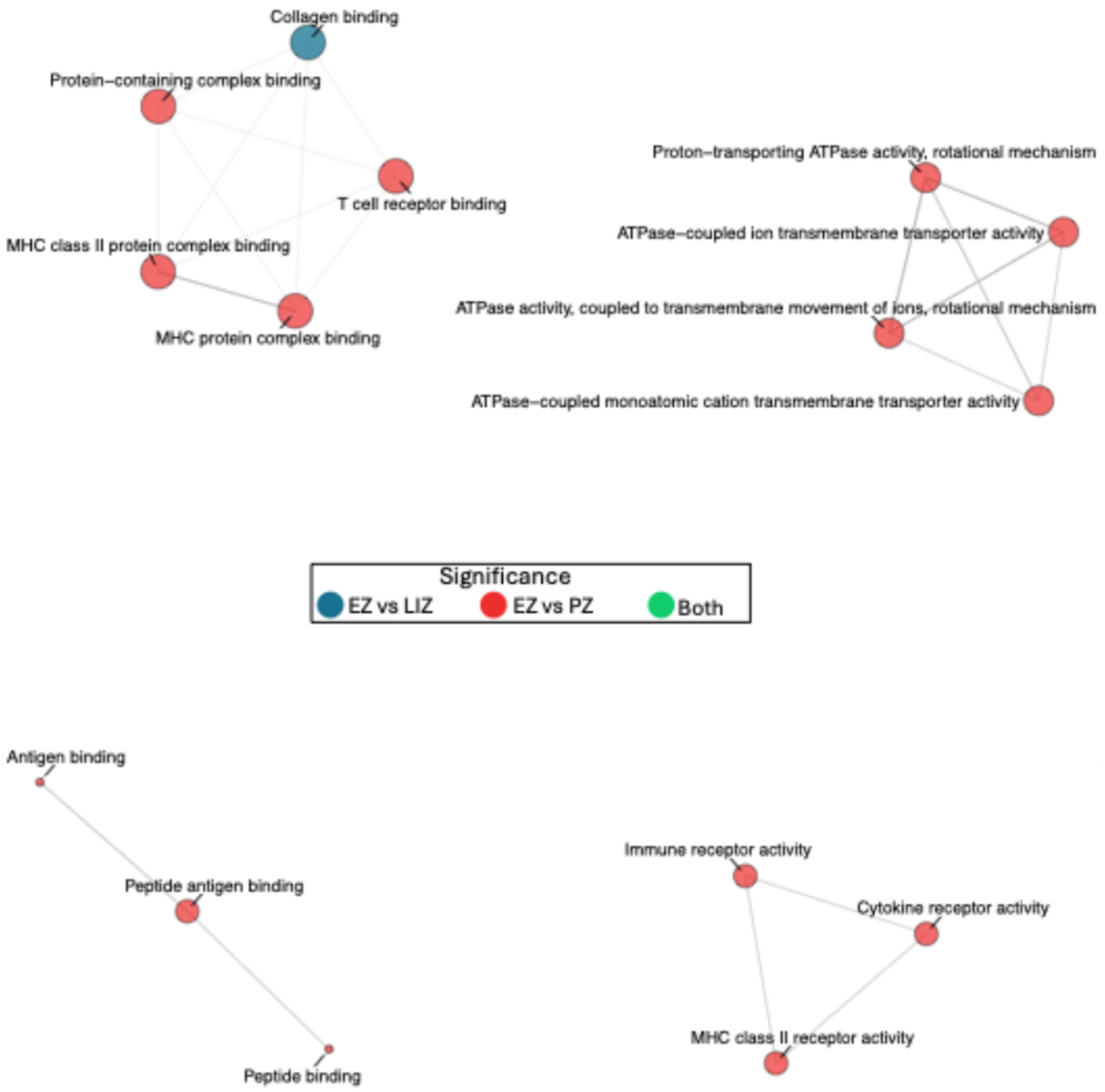


Figure S11 Enrichment map of significant pathways annotated by GO as molecular functions. Nodes represent enriched GO pathways and edges indicate shared genes between the nodes.

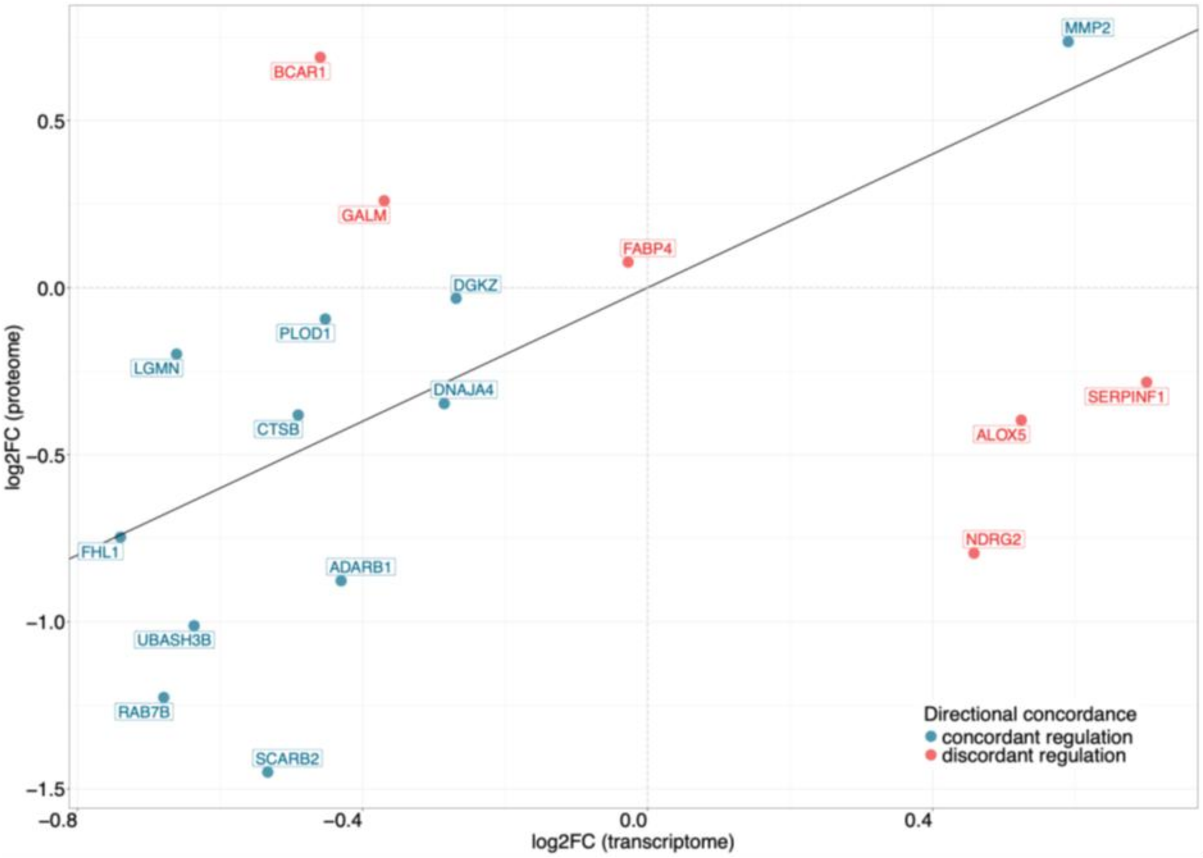


Figure S12 Comparison of transcriptomic and proteomic differential expression in EZ versus LIZ analysis. Each point represents a gene detected in both datasets, plotted according to log2FC in the transcriptome (x-axis) and proteome (y-axis). The diagonal line indicates equal fold changes between transcriptomic and proteomic measurements. Genes showing concordant directional regulation are shown in blue, whereas genes showing discordant directional regulation are shown in red.

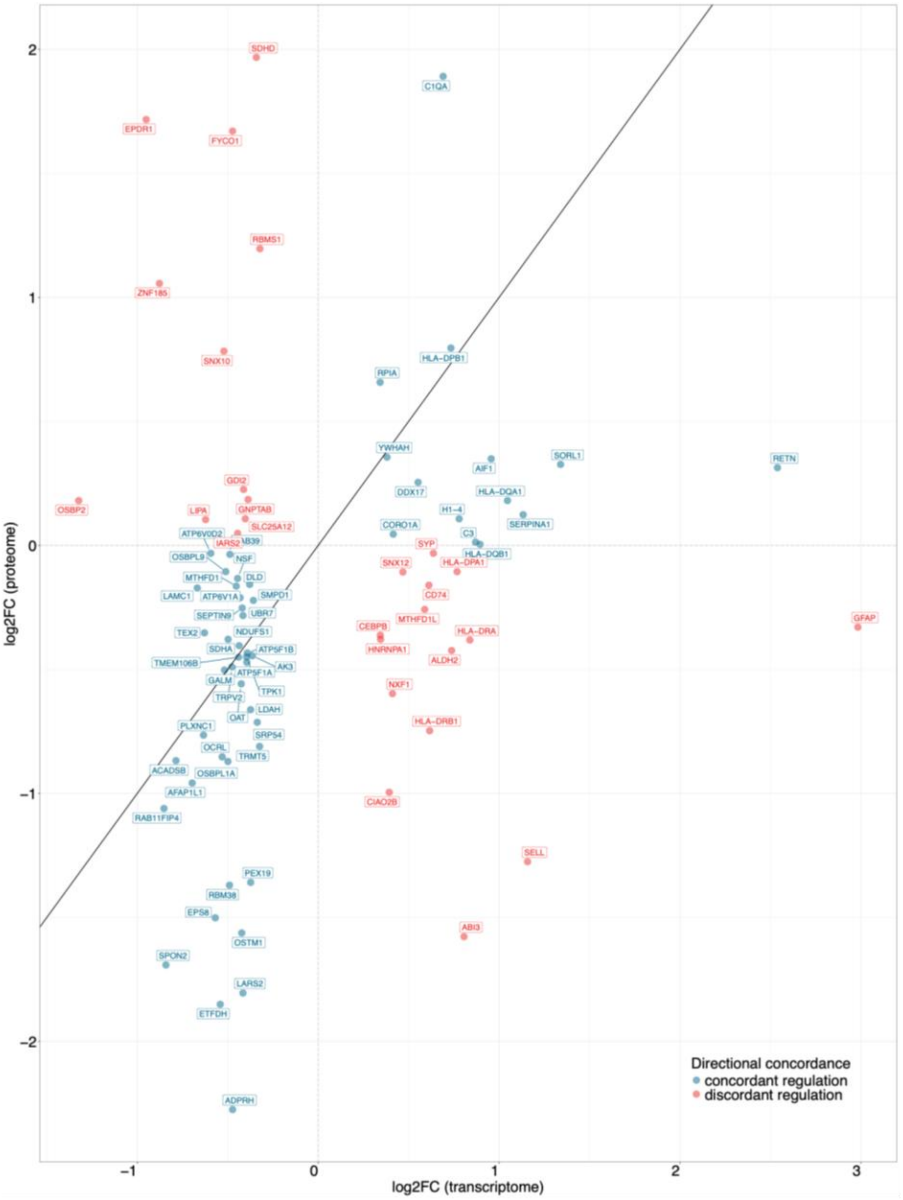


Figure S13 Comparison of transcriptomic and proteomic differential expression effect sizes in EZ versus PZ analysis. Each point represents a gene detected in both datasets, plotted according to log2FC in the transcriptome (x-axis) and proteome (y-axis). The diagonal line indicates equal fold changes between transcriptomic and proteomic measurements. Genes showing concordant directional regulation are shown in blue, whereas genes showing discordant directional regulation are shown in red.
